# *Toxoplasma gondii* serine hydrolases regulate parasite lipid mobilization during growth and replication within the host

**DOI:** 10.1101/2020.11.24.396663

**Authors:** Ouma Onguka, Brett M. Babin, Markus Lakemeyer, Ian T. Foe, Neri Amara, Stephanie M. Terrell, Kenneth M. Lum, Piotr Cieplak, Micah J. Niphakis, Jonathan Z. Long, Matthew Bogyo

**Affiliations:** Department of Pathology, Stanford University School of Medicine, Stanford, CA 94305, USA; Lundbeck La Jolla Research Center, San Diego, CA 92121, USA; Infectious & Inflammatory Disease Center, Sanford Burnham Prebys Medical Discovery Institute, La Jolla, CA 92037, USA; Stanford ChEM-H, Stanford University, Stanford, CA 94305, USA

**Keywords:** α/β-hydrolase superfamily, Serine hydrolases, depalmitoylases, *Toxoplasma gondii*, *Plasmodium falciparum*, Lipid metabolism, CDP-DAG, Phosphatidic acid, Oleic acid, Cholesterol

## Abstract

The intracellular protozoan parasite *Toxoplasma gondii* must scavenge cholesterol and other lipids from the host to facilitate intracellular growth and replication. Enzymes responsible for neutral lipid synthesis have been identified but there is no evidence for enzymes that catalyze lipolysis of cholesterol esters and esterified lipids. Here we characterize several *T. gondii* serine hydrolases with esterase and thioesterase activities that were previously thought to be depalmitoylating enzymes. We find they do not cleave palmitoyl thiol esters but rather hydrolyze short chain lipid esters. Deletion of one of the hydrolases results in alterations in levels of multiple lipids species. We also identify small molecule inhibitors of these hydrolases and show that treatment of parasites results in phenotypic defects reminiscent of parasites exposed to excess cholesterol or oleic acid. Together, these data characterize enzymes necessary for processing lipids critical for infection and highlight the potential for targeting parasite hydrolases for therapeutic applications.

**Highlights:**

- Bioinformatic and biochemical characterization of *T. gondii* serine hydrolases reveals substrate preference between enzymes with similar catalytic fold
- *T. gondii* serine hydrolases previously thought to be depalmitoylases are lipid metabolizing enzymes
- *T. gondii* lipid metabolism pathways utilize enzymes that are viable therapeutic targets

## Introduction

The phylum Apicomplexa contains multiple human parasites, including *Plasmodium falciparum (P. falciparum)* and *Toxoplasma gondii (T. gondii.)*. *T. gondii*, which is an obligate intracellular parasite, asymptomatically infects approximately 30% of the world’s population (Black and Boothroyd, 2000; Boothroyd, 2009; Robert-Gangneux and Darde, 2012). Infection of immuno-compromised individuals such as HIV patients, chemotherapy recipients, and organ transplant recipients can result in toxoplasmosis (Montoya and Liesenfeld, 2004; Tenter et al., 2000). In babies, toxoplasmosis can result in seizures, enlarged liver, enlarged spleen, jaundice, and blindness. Unfortunately, current treatments (Pyrimethamine, Sulfadiazine, Clindamycin, Trimethoprim–sulfamethoxazole) for toxoplasmosis are only effective against acute infections and have side effects that include teratogenic or embryocidal effects, decrease in fertility, and fatal “gasping syndrome” in premature infants (Montoya and Liesenfeld, 2004). For these reasons, a greater understanding of the factors that regulate *T. gondii* asexual lytic cycle has the potential to provide valuable new avenues for therapeutic development for toxoplasmosis.

Previously, our laboratory used a chemical genetic screen to identify a *T. gondii* serine hydrolase *(Tg*PPT1) as a depalmitoylating enzyme (Child et al., 2013; Hall et al., 2011). This approach utilized a small molecule library that contained ~1,200 compounds designed to inhibit cysteine and serine hydrolases (Arastu-Kapur et al., 2008). Inhibition of *Tg*PPT1 resulted in increased host cell invasion as a result of increased parasite motility. However, deletion of the *Tg*PPT1 gene resulted in viable parasites, suggesting that other depalmitoylating enzymes are likely to exist that may be able to compensate for loss of *Tg*PPT1 (Kemp et al., 2013). In further support of this assessment, a homology search by Hidden Markov modeling identified three putative depalmitoylase enzymes in *T. gondii.* These serine hydrolases were named Active Serine Hydrolases *Tg*(ASH) 2, 3 and 4 (*Tg*PPT1 is *Tg*ASH1) and were selected based on the presence of an α/β hydrolase domain and clear sequence similarity to *Tg*PPT1 (Kemp et al., 2013). However, genetic deletion studies provided little insight into the function of these enzymes. While deletion of *Tg*ASH2 or *Tg*ASH3 did not affect growth or infectivity (Kemp et al., 2013), deletion of *Tg*ASH4 caused specific defects in formation of ordered rosettes inside the parasitophorous vacuole, and subsequent inability of parasites to disperse from the site of egress (Foe et al., 2018). In addition, *in vitro* characterization of the purified *Tg*ASH4 enzyme demonstrated that it prefers processing short acyl chains (acetate) suggesting that it is likely not a depalmitoylase. Interestingly, the *Tg*ASH4 deletion phenotype is similar to phenotypes observed when parasites are grown under conditions of excess cholesterol or excess oleic acid (Nolan et al., 2017; Nolan et al., 2018), suggesting that *Tg*ASH4 and possibly *Tg*ASH2 and *Tg*ASH3 might play a role in lipid metabolism in *T. gondii*.

*T. gondii* encodes three genes that produce enzymes involved in neutral lipid synthesis. Two enzymes are responsible for generating cholesterol esters, (acyl-CoA:cholesterol acyltransferases, (*Tg*ACAT1 and 2)) (Lige et al., 2013; Nishikawa et al., 2005; Sonda et al., 2001) and one enzyme generates triacyl-glycerides (acyl-CoA:diacylglycerol acyltransferase (*Tg*DGAT)) (Quittnat et al., 2004), which are both major components of lipid droplets. Deletion of *Tg*DGAT is lethal (Quittnat et al., 2004), and deletion of either *Tg*ACAT1 or 2 results in severe growth defects (Nishikawa et al., 2005) while deletion of both is lethal (Lige et al., 2013; Nishikawa et al., 2005). These results suggest that the pathway for neutral lipid generation is essential for parasite survival. Although the importance of neutral lipid synthesis has been demonstrated for parasite growth and replication, little is known about how parasites utilize scavenged lipids during their intracellular growth.

The *T. gondii* genome contains several genes annotated as putative lipases and carboxy-esterases whose functions have not been experimentally verified. Several or all of these enzymes could be bona-fide acyl-triglyceride lipase(s), even though they lack clear homology with the known mammalian or yeast lipases involved in acyl-triglyceride metabolism. Furthermore, how and when the parasite utilizes scavenged lipids and what enzymes are required for the mobilization of stored neutral lipids is not clear. While it has been proposed that *T. gondii* relies on host enzymes for mobilizing scavenged lipids (Nolan et al., 2017), it remains possible that the parasite has its own internal mechanism for metabolism of these lipids.

In this study, we biochemically characterized the four previously identified ASH α/β–serine hydrolases (*Tg*PPT1 (also known as *Tg*ASH1), *Tg*ASH2, *Tg*ASH3 and *Tg*ASH4) that are conserved across multiple phyla with no known biological function. A BLAST search for conserved motifs identified an acyltransferase motif, a calcium independent phospho-lipase motif, a pentapeptide (GxSxG) lipase motif, and a lipid binding motif. *In vitro* biochemical assays confirm that the enzymes all have both esterase and thioesterase activity with different substrate chain length preferences. Using an LC-MS lipidomics approach, we show that *Tg*ASH4 has a role in parasite lipid metabolism. We also screened and identified inhibitors of recombinant *Tg*ASH2,3 and 4 and show that treatment of parasites phenocopies lipid defects observed in *T. gondii* when grown under excess cholesterol or oleic acid. Together these results define the biochemical properties of the *Tg*ASH 2, 3 and 4 enzymes and suggest that, due to their role in lipid metabolism represent viable targets for treatment of *T. gondii* infections.

## Results

### Sequence analysis and modelling of parasite serine hydrolases

To gain insight into the potential functions of *Tg*ASH proteins, we used sequence alignments and computational structure analyses. *T. gondii* ASH proteins share high protein sequence similarity with members of the α/β-hydrolase superfamily, specifically with serine hydrolases and serine esterases, which include the carboxylesterases, thioesterases, cholinesterases, and lipases (Long and Cravatt, 2011). Sequence alignments confirmed the presence of a conserved catalytic triad (Ser-His-Asp) in each *Tg*ASH protein that can also be found in various homologs in *Plasmodium falciparum* and humans (Figure 1A and S1A). This analysis also revealed the presence of a putative hydrophobic/lipid binding motif (Ghosh et al., 2009; Rajakumari and Daum, 2010) and a calcium independent phospholipase motif (Jenkins et al., 2004) in *Tg*ASH2, *Tg*ASH3, and *Tg*ASH4. In addition, *Tg*AH3 and *Tg*ASH4 each has an additional putative acyltransferase motif (Rajakumari and Daum, 2010).

**Figure 1.**
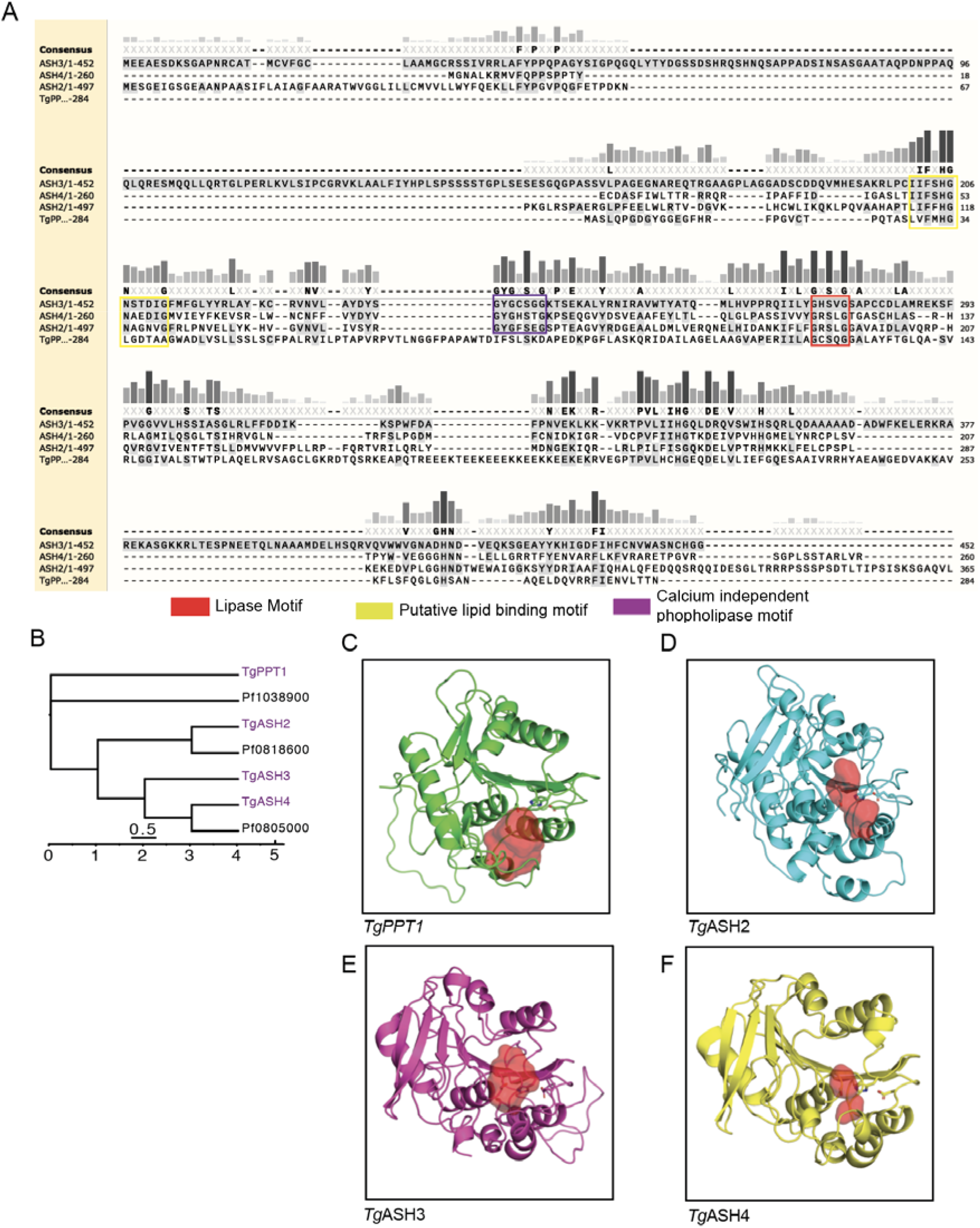
Sequence analysis and modelling of parasite serine hydrolases. (A) Sequence alignment of *Tg*PPT1, *Tg*ASH2, *Tg*ASH3, and *Tg*ASH4 using Clustal Omega. (B) Dendrogram of *T. gondii* and *P. falciparum* serine hydrolases generated using the Clustal Omega algorithm. Branch length represents sequence relatedness. (C) Structural models of (C) *Tg*PPT1 (D) *Tg*ASH2 (E) *Tg*ASH3 and (F) *Tg*ASH4 showing the active site cavity surface (red volume) generated using PyMOL (see also Figure S1).

To place *Tg*ASH proteins in the context of serine hydrolases from humans and the related parasite pathogen *P. falciparum*, we performed phylogenetic analysis of putative serine hydrolase protein sequences. We performed HMMer searches of these SHs against the SwissProt database to generate a list of significant matches (sequence e-value <0.01; SI spreadsheet1) in the human proteome and subsequently performed a multiple alignment of the entire set of hits with the *Toxoplasma*, and *Plasmodium* sequences. This analysis indicated that *Tg*ASH2, *Tg*ASH3, and *Tg*ASH4 are more similar to each other than they are to *Tg*PPT1 (Figure 1B and S1B). *Tg*PPT1 clustered with known acyl-protein thioesterases LYPA1 and LYPA2 (Foe et al., 2015; Kemp et al., 2013), in agreement with our previous studies demonstrating its depalmitoylase activity (Child et al., 2013). By contrast, *Tg*ASH2–4 and the *P. falciparum* serine hydrolases *Pf*0818600 and *Pf*0805000 clustered with the α/β-hydrolase domain containing protein 17 enzymes (ABHD17A, B, and C) and with ABHD13 (ABHDD), all of which are suggested to have depalmitoylase activity. ABHD17s in particular, are thought to be responsible for depalmitoylating N-Ras, thereby regulating the dynamic cycling of Ras localization (Conibear and Davis, 2010). These observations suggested that while *Tg*ASH2–4 also align to human proteins with depalmitoylase activity, their lack of similarity to TgPPT1 suggests they might have distinct substrate preferences.

To gain further insight into the potential unique properties of the parasite serine hydrolases, we generated structural predictions using homology modeling. We aligned parasite proteins to similar proteins for which high resolution crystallographic structures were available. We then selected the best templates chosen using FFAS (Jaroszewski et al., 2011), and generated structural models with MODELER (Sali and Blundell, 1993). For each protein, we built two structures and selected the model that best satisfied the expected geometries of the catalytic triad (Table S1). The final structural models of PPT1, ASH2, ASH3, and ASH4 predict that the catalytic residues all fall within a defined active site pocket (Figures 1C-F, S1C). They also reveal a potential lipid binding domain that overlaps with the acyltransferase motif highlighted in the structures (Ghosh et al., 2009), located near the active site, reinforcing the assignment of the catalytic residues (Figure 1A S1A, Ci-v). Although sequence-based analyses are not sufficient to predict specific enzymatic activity or biological function, in this case they did reveal conserved hydrolase motifs in our *Toxoplasma gondii* serine hydrolases of interest, consistent with our hypothesis that these enzymes share a common catalytic mechanism. Furthermore, phylogenetic clustering suggests that *Tg*ASH2-4 and *Tg*PPT1 have distinct biochemical functions (Figure 1B and S1B). To confirm this hypothesis, the best models for *Tg*ASH2-4 were aligned to *Tg*PPT1using PyMOL “super” (Ho and Gruswitz, 2008). This analysis allows prediction of the size of the active site “cavity and pockets” and correlate that to the size of the substrate that would optimally bind. Based on this structural analysis, we observe that the size/volume of the catalytic cleft is the largest for *Tg*PPT1 (Figure1C), intermediate for *Tg*ASH2 (Figure 1D) and is the smallest for *Tg*ASH3 (Figure 1E) and *Tg*ASH4 (Figure 1F) reflecting possible differences in the sizes of potential substrates that could be accommodated by each enzyme.

### Biochemical analysis of*Tg*ASH esterase and thioesterase activities

Based on homology modeling and the presence of a pentapeptide lipase motif (GxSxG), we hypothesized that *Tg*ASH2-4 are likely to have distinct substrate preferences compared to *Tg*PPT1. In addition, we previously reported that *Tg*PPT1 and *Tg*ASH4 have distinct substrate preferences *in vitro* (Foe et al., 2018), however, the biochemical properties of *Tg*ASH2 and *Tg*ASH3 have not yet been determined. To determine the enzymatic functions of all of the *Tg*ASH proteins, we recombinantly expressed the full-length proteins in *Escherichia coli*, purified them, and subjected them to biochemical analyses.

We first tested the purified enzymes for esterase and thioesterase activities using quenched substrate probes that mimic palmitoylated Ser (QSE) or Cys (QStE) residues (Figure 2A) (Amara et al., 2019; Foe et al., 2018). *Tg*PPT1 showed the highest activity of all the enzymes for these lipid substrates with similar activity for both QSE and QStE. *Tg*ASH2 and *Tg*ASH3 weakly processed both substrates but preferred the thioester substrate. Both enzymes were still six-fold less effective at processing this palmitate mimetic compared to *Tg*PPT1. *Tg*ASH4 showed no activity toward either substrate consistent with its preference for short acyl substrates (Figure 2B-C) (Foe et al., 2018). We next tested whether the enzymes are capable of processing thioesters in general using a substrate with a short acyl chain, S-(4-nitrophenyl)thioacetate (4NPtA) (Figure 2D). In contrast to the palmitoyl substrates, all enzymes were able to process the thioacetate, although *Tg*PPT1 show substantially reduced activity for the acyl substrate compared to ASH2-4, again highlighting the difference in substrate specificity of these serine hydrolases and supporting the hypothesis that only *Tg*PPT1 is a depalmitoylase.

**Figure 2.**
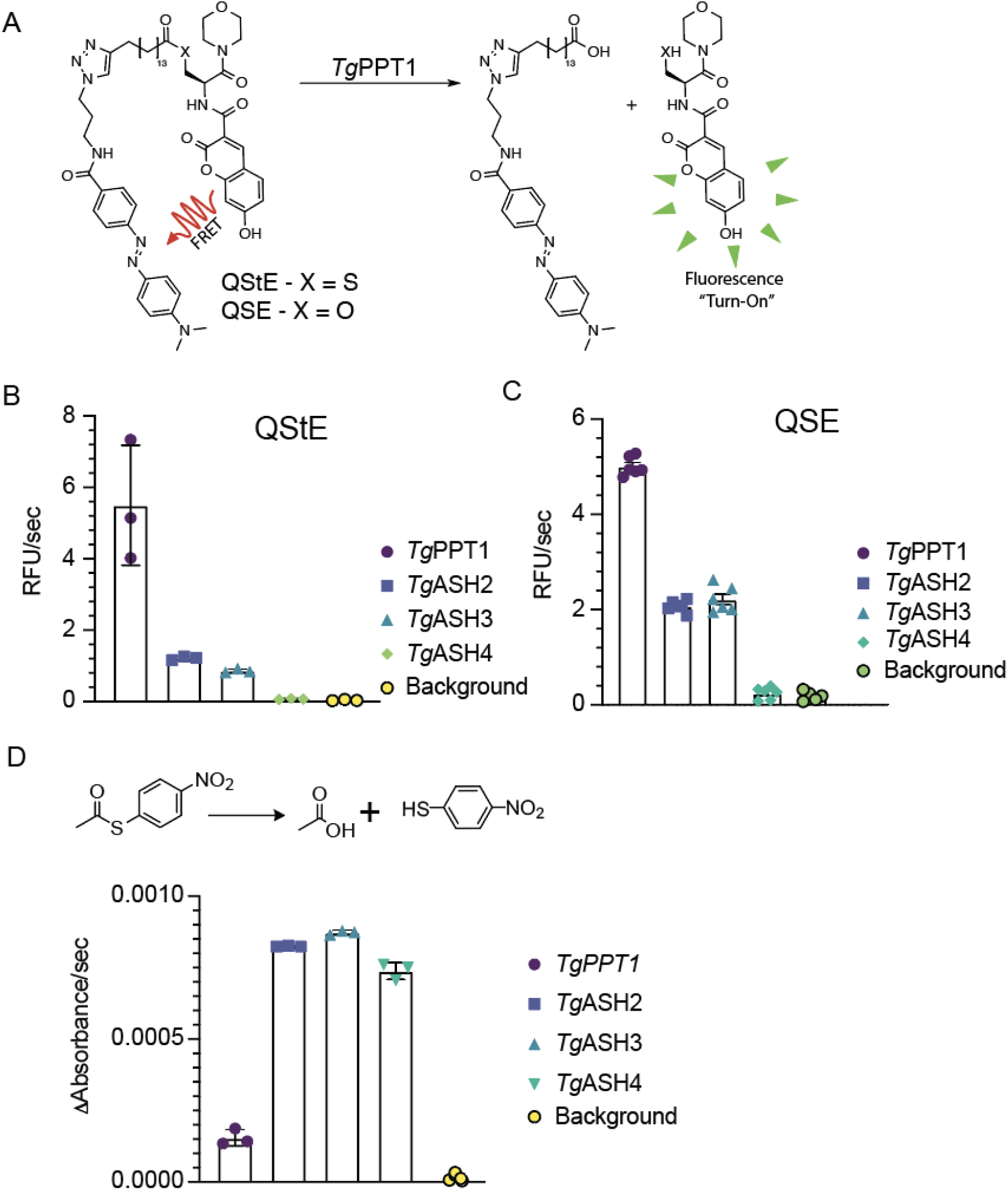
Purified TgASH enzymes exhibit esterase and thioesterase activities. Structures of the Quenched fluorogenic Substrate for Esterase (QSE) and thioEsterase (QStE) activities for measuring depalmitoylase activity. Quantification of the rate of hydrolysis of QStE and (C) QSE by recombinant *Tg*PPT1, *Tg*ASH2, *Tg*ASH3 and *Tg*ASH4 enzymes. Data represents averages of three independent experiments from three technical triplicates. (D) Schematic of hydrolysis of S-(4-nitrophenyl)thioacetate to release the nitro thiophenol reporter (top) and quantification of the rate of hydrolysis by recombinant *Tg*PPT1, *Tg*ASH2, *Tg*ASH3 and *Tg*ASH4. Data represents averages of three independent experiments from three technical triplicates.

To further define the substrate preference for each of the enzymes, we tested a panel of fluorogenic substrates that contain lipids of increasing carbon chain lengths (Figure 3A-C). Consistent with its known function as a depalmitoylase and with its relatively high activity towards QStE, *Tg*PPT1 preferred longer acyl chains (8 and 10 carbons). Interestingly, *Tg*ASH2 was able to process both short acyl substrates and longer lipids but showed a preference for medium length (7 carbons). *Tg*ASH3 and *Tg*ASH4 preferred short acyl chain substrates (2 and 4 carbons) and were unable to process long chain lipid acid esters (Figure 3C and Table 2). These classifications based on substrate specificity also fit with the alignment of the enzymes based on sequence similarities (Figure 1A and B, S1A and B) and homology modeling showing the different size of the active site pockets (Figure 1C-F), suggesting that the enzymes likely have distinct functions *in vivo*.

**Figure 3.**
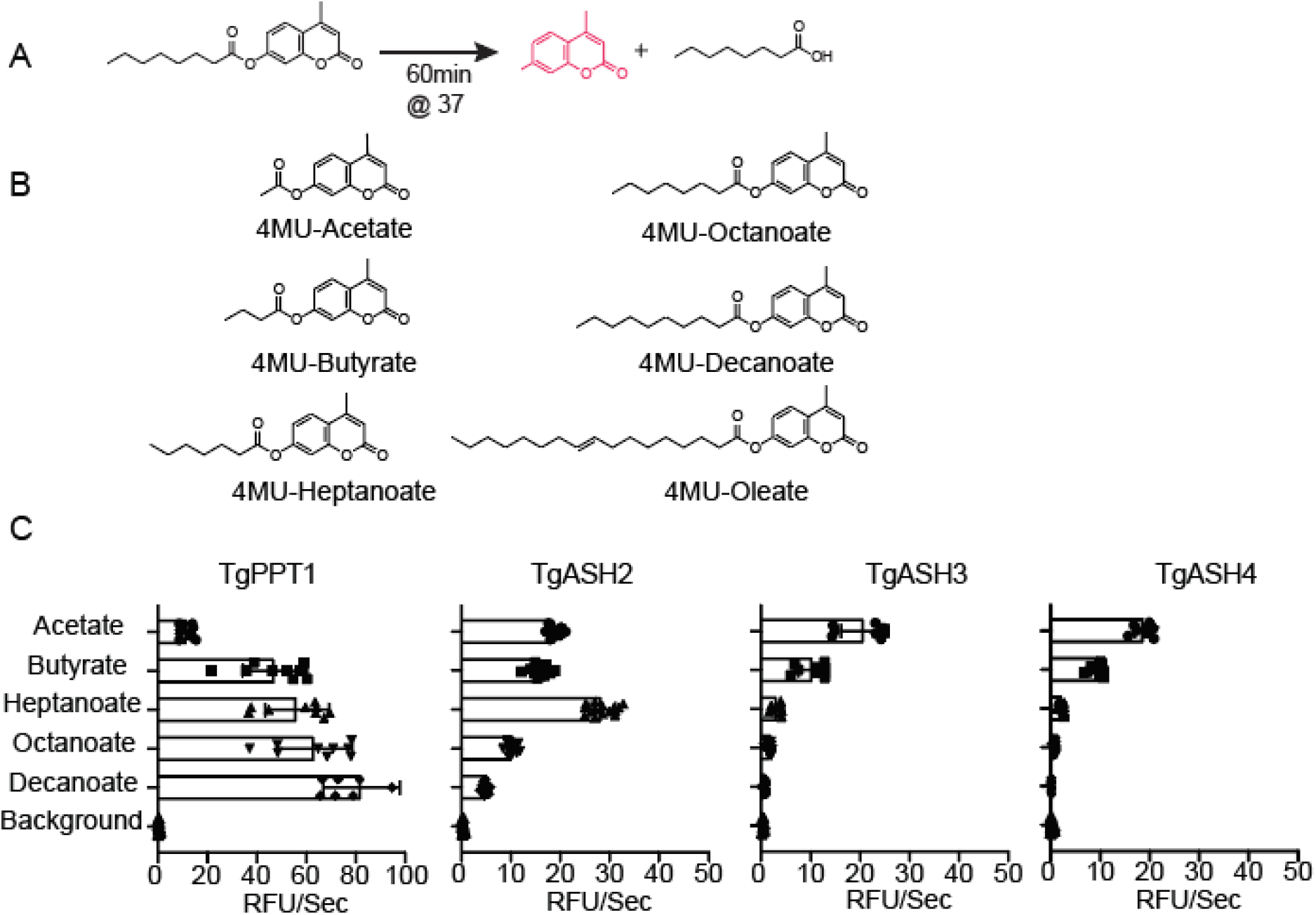
Substrate specificity of TgASH enzymes. (A) General reaction scheme for enzymatic hydrolysis of 4-methylumbelliferone (4MU) substrates. (B) Chemical structures of 4MU substrates with varying lipid chain lengths. (C) Substrate specificity profiles for *Tg*PPT1 and *Tg*ASH2-4 enzymes suing the 4MU substrates. Plots show average turnover rates for each recombinant enzyme as relative fluorescent units. Velocities for each substrate are depicted as relative fluorescence units/sec (λ_ex_ = 365 nm, λ_em_ = 455 nm). Values are the means of triplicates ± standard error.

### *Tg*ASH4 deletion alters the *T. gondii* lipid profile

We previously reported that deletion of *Tg*ASH4 results in parasites that form disordered rosettes (Foe et al., 2018), a phenotype similar to what is observed when parasites are grown in the presence of excess cholesterol or oleic acid (Nolan et al., 2017; Nolan et al., 2018; Robibaro et al., 2002) or when the phosphatidic acid (PA)/diacyl-glycerol (DAG) ratio is dysregulated in *Toxoplasma gondii* (Bullen et al., 2016). Our sequence analysis revealed that *Tg*ASH2-4 each contain a calcium independent phospholipase motif (Figure 1A, S1A, Ci-v), suggesting that these enzymes are likely lipases (Jenkins et al., 2004). Combined with our *in vitro* biochemical data which show that *Tg*ASH2-4 can cleave fatty esters, these observations strongly support a role for all three hydrolases in parasite lipid metabolism (Figure 1A, and SI. 1A). Because deletion of *Tg*ASH2 or *Tg*ASH3 does not result in any phenotypes (Kemp et al., 2013), we believe these enzymes may play redundant or overlapping functions, making genetic studies of function difficult. Therefore, we focused on *Tg*ASH4, which has a clear defect in parasite replication inside the vacuole (Foe et al., 2018). To test our hypothesis that *Tg*ASH4 is involved in lipid metabolism, we performed lipid profiling studies using wild-type and Δ*Tg*Ash4 parasites. We infected primary human foreskin fibroblast (HFF) cells with wild types *T. gondii* parasites as well as parasites in which the TgASH gene was disrupted (Δ*Tg*ASH4). We also generated parasites in the knockout background in which TgASH4 expression was rescued with WT (Δ*Tg*Ash4-ASH4) or catalytically dead (Δ*Tg*Ash4 ASH4-S124A) TgASH4. After 24 h of infection, we collected parasites and extracted lipids for analysis by thin layer chromatography (TLC). We found that loss of *Tg*ASH4 results in accumulation of both phospholipids and neutral lipids compared to wild-type parasites (Figure 4A and B). This accumulation was rescued by expression of wild type (Δ*Tg*Ash4-ASH4) but not catalytically dead mutant (Figure 4A and B).

**Figure 4.**
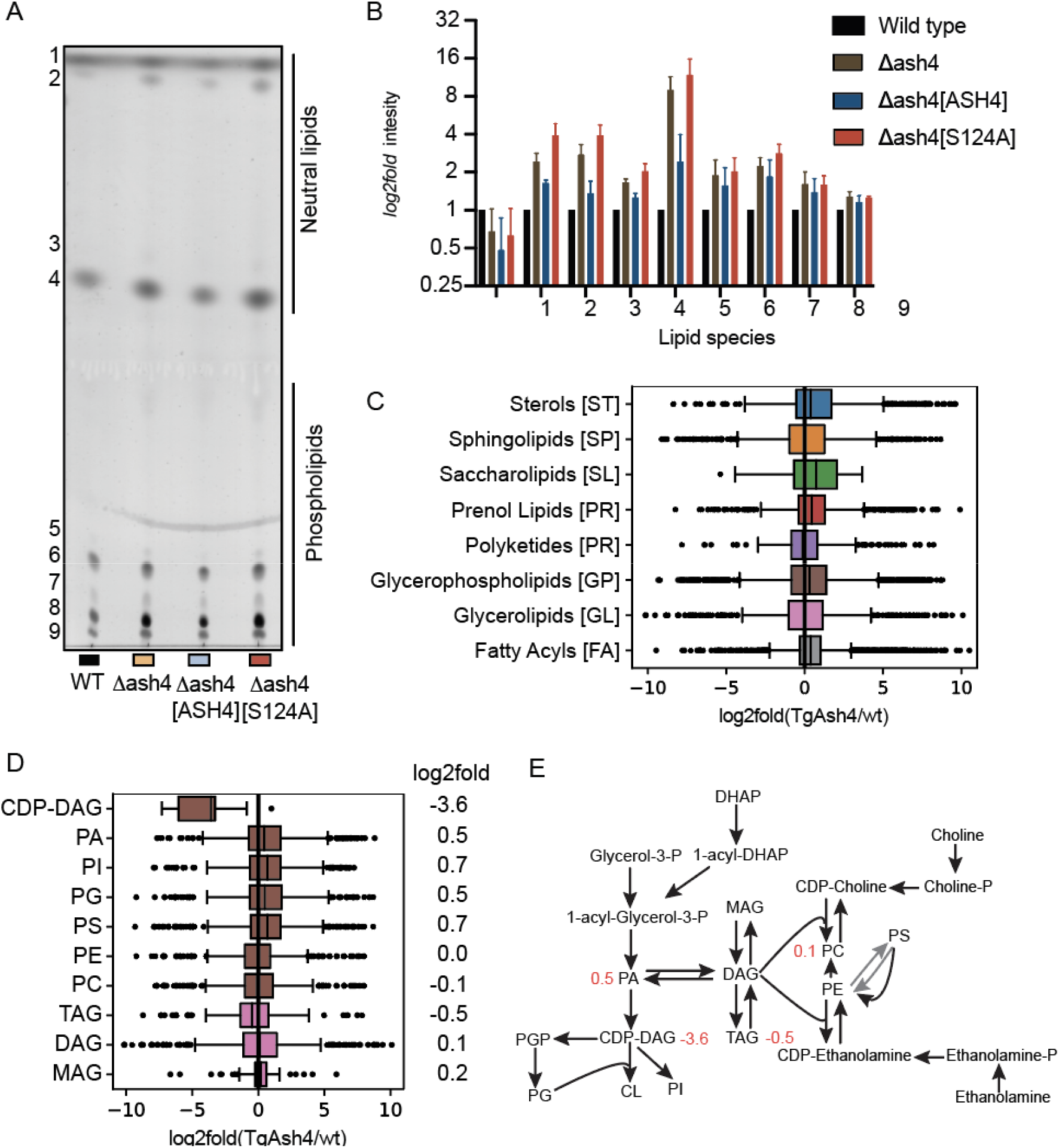
*Tg*ASH4 deletion alters the *T. gondii* lipid profile. (A) Image of TLC separation of lipids extracted from wild type, Δ*Tg*Ash4, Δ*Tg*Ash[ASH4] and Δ*Tg*Ash4 *Tg*ASH4 [S124A] parasites. Lipids were stained by cupric sulfate. (B) Quantification of stained lipids from TLC analysis and plotted as a log2fold change. Box plots showing lipid distribution in Δ*Tg*Ash relative to wild type parasites grouped by (C) main general categories of lipids and (D) the sub-class of glycerophospholipids with log2fold changes indicated. (E) schematic of glycerolipid metabolism pathways, with numbers in red indicating log2fold change in Δ*Tg*Ash4 vs wild type. Abbreviations: Cytidine-diphosphate-diacylglycerol (CDP-DAG), Phosphatidic acid (PA), Phosphoinositol (PI), Phosphatidyl-glycerol (PG), Phosphatidylserine (PS), Phosphatidyl ethanolamine (PE), Phosphatidyl choline (PC), Triacylglycerol (TAG), Diacylglycerol (DAG), Monoacylglycerol (MAG), Glycerol 3-phosphate (Glycerol-3-P), Phosphatidyl-glycerol-phosphate (PGP), Cardiolipin (CL), Dihydroxyacetone-phosphate (DHAP), Cytidine-diphosphate-choline (CDP-choline), Cytidine-diphosphate-ethanolamine (CDP-Ethanolamine) (see also Figure S2).

To determine the identity of the lipids that accumulated in the absence of *Tg*ASH4, we extracted lipids from wild-type and Δ*Tg*Ash4 parasites and analyzed the total lipidome by LC-MS. To increase the number of lipids identified, we used both positive and negative electrospray ionization (ESI) modes for MS analysis. In general, we observe distinct lipid profiles in the parasites lacking *Tg*ASH4 compared to wild type parasites (Figure. 4C-D and S2A, B), supporting our hypothesis that *Tg*ASH4 plays a role in lipid metabolism in *T. gondii*. A more detailed survey of the lipidomic data confirmed general defects in glycerolipid metabolism (Figure 4D–E, S2 B). Specifically, we found that levels of cytidine diphosphate-diacylglycerol (CDP-DAG) lipids were on average 16-fold decreased in the knock out parasites. We also observed a slight increase is phosphatidic acid (PA) upon loss of TgASH4 expression. Both of these classes of lipids are central intermediates in phospholipid and glycerophospholipid metabolism (Figure 4D, E and S 2B). Synthesis of PA is the initiating event in phospholipid biosynthesis in both prokaryotic and eukaryotic organisms (Henry et al., 2012). PA is then converted into CDP-DAG, the central liponucleotide intermediate for phospholipid biosynthesis (Pascual and Carman, 2013). In eukaryotes, PA is a precursor for both CDP-DAG and diacylglycerol (DAG). CDP-DAG is used to make phosphoinositol (PI), phosphatidylglycerol (PG) and cardiolipin (CL), while DAG is required for phosphatidylcholine (PC), phosphatidylethanolamine (PE), and triacylglycerol (TAG) synthesis (Carman and Han, 2019; Henry et al., 2012) (Figure 2E).

### Identification of small molecules that inhibit *Tg*ASH proteins

Given the likely roles for *Tg*ASH2-4 in lipid metabolism, we sought to identify inhibitors of these enzymes that could be used to further explore their biological function during *T. gondii* infection. Our laboratory has compiled a library of small molecules that covalently react with cysteine and serine hydrolases. This library contains around 1,200 compounds that have diverse scaffolds attached to reactive electrophiles (Arastu-Kapur et al., 2008; Child et al., 2013; Ponder et al., 2011). We therefore performed our screening against the *Tg*ASH proteins using the set of approximately 400 compounds that are likely to target serine hydrolases. To identify inhibitors, we pooled the recombinant *Tg*ASH enzymes and performed a gel-based competitive activity-based profiling assay (Figure S3). We pretreated the mixture of enzymes with each compound for 30 minutes, after which, we labeled the enzyme mixture with the broad-spectrum serine hydrolase probe, FP-TAMRA (Figure S3A, B). Compound binding appeared as a loss of labeling by the FP-TAMRA probe (Figure S4, S5). Because the enzymes have distinct molecular weights they can be resolved using SDS-PAGE, we could quantify the potency and selectivity of each compound for each enzyme target in a single lane of a gel.

As an initial cutoff, we selected compounds that reduced FP-TAMRA labeling by at least 60% at 33 μM (Figure 5A). To validate the hits, we used a secondary assay to measure their ability to inhibit processing of the fluorogenic 4MU-octanoate substrate (Figure 5B and Figure SI 6B). In this assay, we screened the enzymes separately at lower concentrations of inhibitors. Confirmed hit molecules that were active in both assays shared a core chloroisocoumarin scaffold (JCP341,JCP342,JCP343, JCP348, and JCP388: Figure 5B). Chloroisocoumarins are electrophilic traps that form irreversible covalent bonds with active site nucleophiles of serine hydroalases. The compounds are initially attacked by the active site serine residue to form a reversible ester linkage and then by subsequent irreversible covalent modification by a nearby histidine. We validated inhibitors by calculating apparent IC_50_ values following dose dependent inhibition studies using a fixed time point assay (Figure S6B). Most of the screening hits inhibited all three serine hydrolases, although with overall differences in potency (IC_50_) and maximum level of inhibition for each hydrolases. However, two of the lead compounds from the screen, JCP343 and JCP348, were potent inhibitors of ASH2-4 but showed virtually no activity for *Tg*PPT1, making them highly valuable for studies of *Tg*ASH2-4 function.

**Figure 5.**
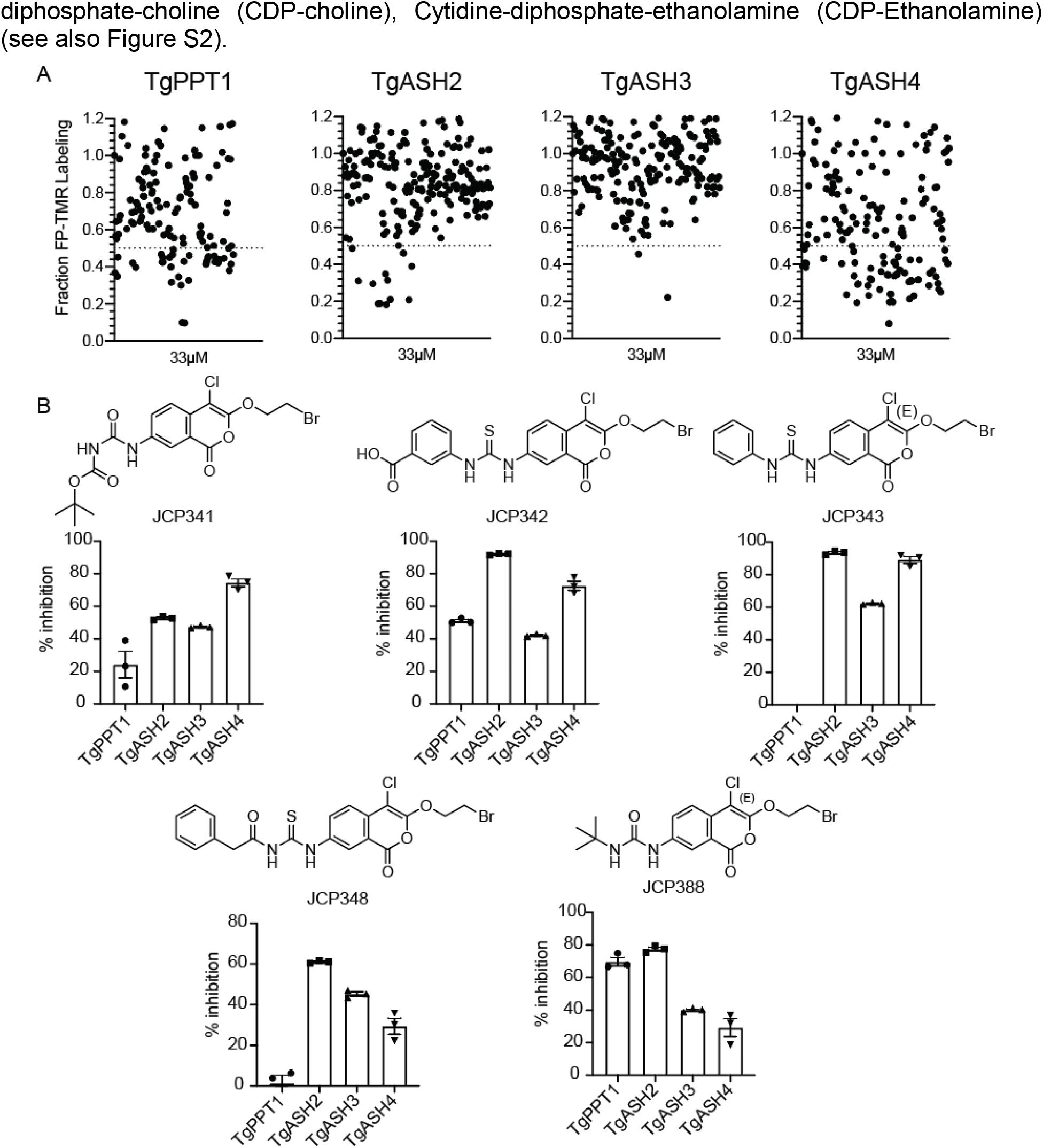
Identification of small molecules that inhibit *Tg*ASH proteins. (A) Dot plot of screening results of the gel-based competition assay with FP TAMRA and serine reactive libraries against *Tg*PPT1, *Tg*ASH2, *Tg*ASH3, and *Tg*ASH4. The enzymes were pre-incubated with 33μM compound, or DMSO. Results display the residual TAMRA signal, normalized to the DMSO control and plotted as a percentage of TAMRA labeling relative to the DMSO control. (B) Structures of selected compounds and their percent inhibition of respective enzymes at 10μM as assayed by turnover of 4MU-octanoate (*Tg*PPT1, *Tg*ASH2 and *Tg*ASH3) or gel-based competitive FP-TAMRA labeling for *Tg*ASH4 (see also Figure S3, S4 and S5).

### Inhibition of TgASH proteins impairs parasite infection

For *T. gondii* to successfully establish an infection, the parasite must invade host cells (Carruthers and Boothroyd, 2007), undergo asexual replication (Clough and Frickel, 2017), egress (exit) from host cells, and invade new cells (Black and Boothroyd, 2000; Blackman and Carruthers, 2013). Therefore, to examine whether the inhibition of *Tg*ASH proteins impacts any of the asexual lytic stages of *T. gondii*, we treated extracellular tachyzoite parasites with the newly identified inhibitors of *Tg*ASH2-4. We used plaque and replication assays to measure parasite fitness upon compound treatment. Plaque assays can be used to assess the overall viability of the parasites through multiple rounds of the asexual lytic cycle. In these studies, we included the FDA-approved drug, orlistat, as a positive control, since it inhibits fatty acid synthase and has been shown to block lipid metabolism processes in humans and in *Plasmodium falciparum (Yoo et al., 2020)*. All inhibitors severely impaired parasite growth with 50–75% reduction in plaque numbers compared to a DMSO control (Figure 6A, B, and Figure S6C). These data strongly suggest that TgASH2-4 are important for parasite intracellular growth.

**Figure 6.**
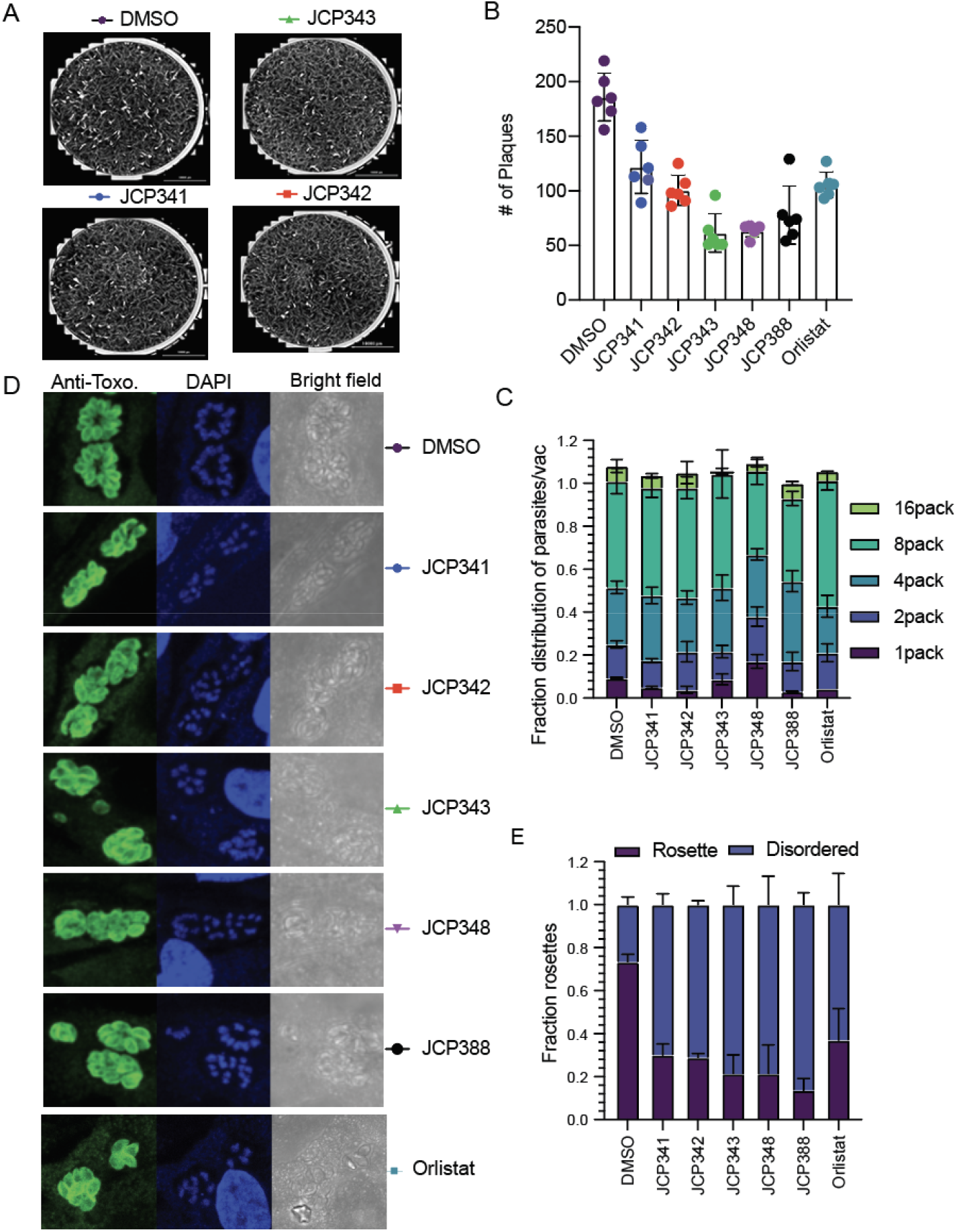
Inhibition of TgASH proteins impairs parasite infection. (A) Representative images of plaques formed by parasites after pretreatment of the parasites with either DMSO or the indicated screening hits (10 μM final). (B) Quantification of total plaques numbers upon compound treatment. Data represent averages (bar plot) from three independent experiments performed in technical triplicates. (C) Quantification of fractions of parasites per vacuole of parasites treated with either DMSO or selected inhibitors. (D) Indirect immunofluorescence images of wild-type parasites treated with either DMSO or indicated inhibitors. Parasites are stained with an anti-Toxo antibody (green) and nuclei of host cells stained with DAPI (blue). (E) Quantification of fractions of parasites in ordered and disordered rosettes for DMSO or compound treated wild type parasites. The graph presents averaged results from three independent experiments performed in technical triplicate. For replication assays and rosette assay, at least 100 vacuoles/per condition were counted (see also Figure S6).

To investigate which stage of the asexual lytic cycle is impacted by *Tg*ASH2-4 inhibition, we first analyzed the rate of replication in either the presence or absence of inhibitors. The rate of parasite replication can be determined by quantifying the number of parasites per vacuole upon treatment with either DMSO or inhibitor (see methods for further information). Parasites pretreated with the identified hits resulted in significantly slower growth rate (less 8 and 16 parasites/vacuole) compared to DMSO treated controls, suggesting a disruption in the rate of parasite replication (Figure 6C). The relative potencies of the compounds in this assay matched the patterns we observed in the plaque assay (Figure 6B) and the potencies for the recombinant *Tg*ASH proteins (Figure 5B).

*T. gondii* tachyzoites divide asexually in the parasitophorous vacuole (PV) through a process called endodyogeny. During this process, parasites sequentially form daughters that are connected by a residual body, resulting in the formation of a flower shaped “rosette” of daughter parasites. We previously showed that deletion of *Tg*ASH4 results in disordered rosettes in which cell division is disrupted and daughter pairs have individual residual bodies with altered parasite morphologies. We analyzed the effect of the ASH inhibitors from our library screen on the formation of rosettes inside the host cell and found that treatment of parasites with a 10 μM dose of the *Tg*ASH2-4 inhibitors resulted in a loss of normal rosettes and formation of the same type of disordered rosettes that we observed in the *Tg*ASH4 knock out parasites (Figure 6D, E). This observation suggests that *Tg*ASH2-4 inhibitors, like the general inhibitor of lipid metabolism, orlistat, produce a phenotype that we previously observed for the *Tg*ASH4 deletion mutant and which mimics the effects of excess cholesterol or oleic acid (Foe et al., 2018).

## Discussion

Although it is possible to predict protein function by assessment of sequence similarity to proteins with known functions (Dolinski and Botstein, 2007; Train et al., 2017), it is difficult to identify proteins with similar function if they lack sequence similarities. The α/β-hydrolase fold superfamily of enzymes comprises enzymes that contain a conserved catalytic triad (a serine nucleophile, catalytic acid and histidine) and common structural fold. Despite conserved active sites and structural features, α/β-hydrolases perform diverse biological functions (Carr and Ollis, 2009; Ollis et al., 1992). Thus, the α/β-hydrolase superfamily is an excellent model to begin to understand the structure–function relationships of a common structural protein fold (Long and Cravatt, 2011). In this work, we employed bioinformatic, biochemical, cell biological, and pharmacological approaches to define the functions of genetically related serine hydrolase enzymes in the important human parasite pathogen *T. gondii*.

Sequence and structural analyses identified protein domains conserved among all four enzymes. It also highlighted some key differences, including a conserved calcium independent phospholipase motif (Jenkins et al., 2004; Rajakumari and Daum, 2010) that is lacking in *Tg*PPT1 and the presence of putative acyl transferase motif H(x)4D (Rajakumari and Daum, 2010) in *Tg*ASH3 and *Tg*ASH4. Although all of the ASH proteins aligned with human proteins annotated as depalmitoylases, our biochemical analyses indicate that only *Tg*PPT1 is likely a *bona fide* depalmitoylase, while *Tg*ASH2, *Tg*ASH3, and *Tg*ASH4 exhibit only weak or no activity towards long chain thioesters. Together these biochemical results suggest that ASH proteins should be classified into three subsets based on their enzymatic activities: (1) depalmitoylases (i.e. long chain lipid thioesters) – *Tg*PPT1; (2) medium chain lipid esterases – *Tg*ASH2, and (3) short chain lipid esterases/deacylases – *Tg*ASH3 and *Tg*ASH4. These classifications based on substrate specificity also fit with the alignment of the enzymes based on sequence homology (Figure 1B) and suggest they likely have distinct functions *in vivo*. While conserved positions in α/β-hydrolases define properties that are shared within the entire family (for example, the catalytic triad), they do not fully explain functional diversity or biological function (Long and Cravatt, 2011). These findings emphasize the complementary nature of bioinformatic and biochemical investigations. Whereas we were unable to predict substrate preference based on sequence similarity and homology modeling alone, clustering analysis suggested that the enzymes differed with respect to conserved domains, prompting our initial biochemical hypotheses.

Deletion of *Tg*ASH4 yields parasites that form smaller plaques, disordered rosettes, and dispersion defects compared to wild-type parasites (Foe et al., 2018). In addition, the balance between DAG and PA is important for *T. gondii* growth and replication. Furthermore, inhibition of enzymes involved in PA and DAG formation results in defects in rosette formation (Bullen et al., 2016). Our data supports a model in which these defects are caused by disruption of lipid metabolism. The lipid profiles of parasites lacking *Tg*ASH4, or Δ*Tg*ASH4 rescued with wild type*Tg*ASH4 or Catalytic dead *Tg*ASH4 accumulate both phospholipids and neutral lipids. Specifically, parasites lacking *Tg*ASH4 have lower levels of cytidine diphosphate diacylglycerols (CDP-DAG) and an increase in phosphatidic acids (PA) and several sterols. CDP-DAG and PA are essential lipid intermediates that are involved in *de novo* synthesis of all phospholipids in both eukaryotes and prokaryotes. CDP-DAG synthase (CDS) produces CDP-DAG from PA and cytidine triphosphate. *T. gondii* contains two phylogenetically divergent CDS enzymes (Cds1 and Cds2) that reside in the ER and the apicoplast, respectively (Kong et al., 2017). Both *Tg*Cds1 and *Tg*Cds2 are essential and conditional knockdown results in parasites with reduced growth rates that could not be restored by addition of exogenous lipids, suggesting that *de novo* synthesis of lipid intermediates is essential to parasite growth (Kong et al., 2017). These observations highlight the critical link between coordinated regulation of lipid flux and cell division.

Our prior studies showed that Δ*Tg*ASH4 parasites tend to have distended residual bodies that are derived from a single parasite and multiple cell division defects. These phenotypes include incomplete cytokinesis at the apical end after division has initiated, incorrect initiation of division from the basal side, defects in endodyogeny, and defects in cytokinesis (Foe et al., 2018). Our current pharmacological experiments yielded complementary observations, wherein parasites treated with the *Tg*ASH2-4 inhibitors exhibited growth and replication defects. Importantly, these phenotypes are similar to those observed for parasites grown under excess cholesterol or oleic acid. In addition, treatment of parasites with Orlistat, a drug that is known to target lipid pathways (Ross and Fidock, 2019; Yoo et al., 2020), also results in disordered rosettes. These results suggest that *Tg*ASH2-4 inhibitors are generally involved in parasite lipid metabolism.

In summary, we have characterized a set of serine hydrolases that were previously thought to be depalmitoylases based on sequence alignment, but our data suggest that they are in fact esterases involved in parasite lipid metabolism. We show that parasites lacking *Tg*ASH4 have dysregulated lipid metabolism resulting in slight increase in PA and a substantial decrease in CDP-DAG, two key intermediates in the generation of phospholipid biogenesis that supply the entire endomembrane system with phospholipids (Carman and Han, 2019). In addition, FA, cholesterol esters, and sphingolipids are increased in the absence of *Tg*ASH4. Although we cannot yet identify the particular metabolic pathway that links *Tg*ASH4 activity to lipid disruption, our data support a model in which either *Tg*ASH4 is either directly or indirectly involved in CDP-DAG synthesis. A balance of PA/DAG is important for lipid metabolism and cell physiology since both act as precursors for other phospholipids and as signaling molecules. An abnormal increase in lipid levels results in cells susceptible to fatty acid induced lipotoxicity, defects in vacuolar fusion and acidification, autophagy, and cell wall integrity. In the future, we plan to perform further analysis of higher order combinations of deletions of the *Tg*ASH proteins in order to more specifically characterize the lipid metabolism pathways in which they operate.

### Significance

The intracellular protozoan parasite *Toxoplasma gondii* must scavenge cholesterol and other lipids from the host to facilitate its intracellular growth and replication, while avoiding excess accumulation and lipotoxicity. However, when and how parasites mobilize scavenged lipids required for growth is still an open question. We have bioinformatically and biochemically characterized several *T. gondii* serine hydrolases that are conserved across multiple phyla with no known function. We have shown that these enzymes, although initially thought to be depalmitoylases, are rather metabolic enzymes important for lipid regulation within the parasite during growth and development. The lipidomic and pharmacological studies presented here also provide further support for the essential role of lipid metabolism in *T. gondii* and suggest that targeting the *Tg*ASH proteins could be a viable strategy for treatment of *T. gondii* infections.

## Materials and methods

### Parasite culture

Parasites were grown in human foreskin fibroblasts (HFFs) using a mixture of Dulbecco’s modified Eagle’s medium (DMEM) supplemented with 10% FetalPlex animal serum complex (Gemini Biotech, catalog no. 100602), 2mM L-glutamine, and a cocktail of 100μg/ml penicillin, and 100 μg/ml streptomycin. Parasites were cultured at 37°C in 5% CO_2_.

### Plasmid construction

Bacterial expression construct was created by PCR amplification of the *ASH4* open reading frame (ORF) from the strain RH cDNA using primers. The PCR product was digested with Nde1 and BamH1 and ligated into pET-28A, resulting in 6XHis-appended to the N-terminus of *Tg*PPT1, *Tg*ASH2, *Tg*ASH3 and *Tg*ASH4.

### Phylogenetic analysis

Sequence alignments were performed using Clustal omega algorithm (Sievers and Higgins, 2018). For structural modeling of the proteins, we searched for the best structural template by performing sequence alignment of a target sequence with the sequences of proteins, for which experimental structure is known. After which we used MODELER program (Ollis et al., 1992), this program builds three-dimensional model that satisfy various spatial restraints of the template structure. FFAS program (Jaroszewski et al., 2005) was then used to find the best templates (Jaroszewski et al., 2011). Building the final structural models has been done by MODELER program (Sali and Blundell, 1993). For each protein three structures were build, and only a single final model that satisfy/model best the geometries of the catalytic triad SDH in the active site of the enzymes and its FFAS score is the lowest has been selected for further analysis for each enzyme. The information about templates used for model building, including sequence identity, FFAS scores, positions of catalytic residues in the model and brief characteristics of the models used for final selection of the models is summarized in Table S1. The final models selected for structural analysis have been built using the following templates from PDB: for ASH1 – 5syn_A, ASH2 – 6imp_A, ASH3 – 5g59_A and for ASH4 – 5g59_A. The best models for ASH2-4 were aligned (PyMOL “super”) to that of ASH1 (*Tg*PPT1). The region of the active site cavity was visualized in using the “Cavity and Pockets” surfaces generated by PyMOL, taking into consideration the contiguity of the surfaces as well as the direction of the serine nucleophile. HOLLOW^1^ (1.1) was used to generate a cast of the protein surface in a 15.0 Å radius centered on the active site serine oxygen (OG), with interior probe and surface probe radii set to 1.4 and 4.2 Å respectively (Ho and Gruswitz, 2008). The resulting casts were manually trimmed to coincide with the region of interest observed in PyMOL.

### Protein purification

HIS6-PPT1 (rPPT1), HIS6-ASH2 (rASH2), HIS6-ASH3 (rASH3) HIS6-ASH4 (rASH4) and with their mutant counterparts HIS6-PPT1S128A, (rPPT1S128A), HIS6-ASH2S192A (rASH2S192A), HIS6-ASH3S277A (rASH3S192A), and HIS6-ASH4S124A (rASH4S124A) were expressed from the pIF22 and pIF22-S124A plasmids in BL21-CodonPlus (DE3)-RIL *Escherichia coli* (Agilent Technologies, catalog no. 230245). Expression was induced with 0.5 mM IPTG (isopropyl-B-D-thiogalactopyranoside) for 16 h at 19°C. Recombinant proteins were purified as previously described(Child et al., 2013; Foe et al., 2015; Foe et al., 2018; Yoo et al., 2020) with the following modifications: 0.02% NP-40 detergent and 10% glycerol were included in both the lysis and wash buffer. Protein concentrations were quantitated by bicinchoninic acid (BCA) assay.

### *In vitro* biochemical assays

4MU substrate assays were performed as previously described(Foe et al., 2018). Substrates were used at a final concentration of 10 μM. Enzymes were used at a 50nM final concentration for the experiments represented in Fig. 1A. Enzymes were used at a final concentration of 50 nM for the experiments represented in Fig. S1B in the supplemental material. Fluorescence (Ex = 335 nm and Em = 450 nm) was measured every minute on a Cytation 3 imaging reader (BioTek, Winooski, VT, USA) for 60 min. 4-Nitrophenol acetate (4NPA) and 4-nitrophenol thiol-acetate (4-S-NPA) assays were performed as previously described for 4-nitrophenol octanoate (4NPO) (Foe et al., 2018). Enzymes were used at a final concentration of 150 nM. Absorbance was monitored on a Cytation 3 imaging reader (BioTek, Winooski, VT, USA).

### In vitro depalmitoylase activity assays

QStE and QSE assays were performed as previously described method (Amara et al., 2019; Foe et al., 2018). Purified enzymes were diluted in reaction buffer (20 mM HEPES, 150 mM NaCl, 10 mM CHAPS, pH 7.4) to a final concentration of 150μM, and pipetted into each well in a 384-well black plate. Fluorogenic substrates were then added at a final concentration of 20 μM. Reactions were incubated at 37°C and fluorescence was measured (Ex = 410 nm, Em = 450 nm) over a period of 60 minutes on a Cytation 3 imaging reader (BioTek, Winooski, VT, USA). Reaction rates were determined by linear regression analysis of the initial linear phase of hydrolysis. Experiments were performed in three technical replicates and three biological replicates.

### Steady-state kinetics

Parameters of steady state kinetics were determined by measuring the increase of fluorescence of hydrolyzed fluorogenic substrates (4MU (Foe et al., 2018)). Stock solutions of fluorogenic substrates were diluted in reaction buffer (1X PBS, 0.05% Tx-100, pH 7.4) and pipetted into 384-well black plates, at a series of concentrations ranging from 0.8 μM to 0.391 μM. Recombinant enzymes were then added to a final concentration between 50 nM. Reaction progress was recorded (Ex = 335 nm, Em = 450 nm) over a period of 30-60 minutes at 37°C on a Cytation-3 imaging reader (BioTek, Winooski, VT, USA). Concentrations of hydrolyzed products were determined using a standard concentration curve (unconjugated 4MU). Reaction rates were determined by linear regression analysis of the initial velocity (V0) of hydrolysis. Experiments were performed in triplicates. The kinetic constants, Km and Kcat, were obtained by non-linear regression curve fit of Michaelis-Menten parameters; V_0_=[E]*K_cat_*[S]/(K_m_+[S]), V_0_ (initial reaction rate), E (enzyme concentration), S (substrate concentration), K_m_ (Michaelis constant) and K_cat_ (turnover number) to these data using GraphPad Prism. Catalytic efficiencies were calculated from the product of K_cat_/K_m_.

### FP-TAMRA competition assays

For competition assays, 300 ng of purified enzyme was incubated with DMSO or inhibitors at 10 mM for 30 minutes on ice, then 1 mM of FP-TAMRA was added and samples were incubated for additional 30 at 37°C. Reactions were quenched with reducing SDS sample buffer, and the entire sample was resolved by SDS-PAGE. A typhoon flat-bed scanner was used to scan the gel (532-nm laser, 610-nm filter, PMT800).

### Plaque assays

Confluent monolayers of HFFs grown in 6 well plates were infected with freshly egressed tachyzoites isolated from host cells by syringe lysis and filtered with a 5-μm filter to remove host cell debris. The number of parasites per microliter was counted on hemocytometer. A total of 200 parasites were added to confluent HFFs in 6-well dishes. Parasites were grown for 7 days, fixed with cold methanol, and stained with crystal violet.

### Immunofluorescence microscopy

Parasites were allowed to infect confluent HFFs on coverslips for 24 h, after which coverslips were fixed 4% paraformaldehyde. Cover slips were permeabilized with 0.2% Triton X-100 in 1X phosphate-buffered saline (PBS) and blocked in 3% bovine serum albumin (BSA) in 1X PBS for 30 min. Parasites were stained with anti-Toxo-fluorescein isothiocyanate (anti-Toxo-FITC) (Thermo Fisher, catalog no. PA1-7253) antibody by incubating overnight at 4°C in 3% BSA in 1X PBS at 1:1000 dilution. After incubation with primary antibodies, coverslips were washed 3 times with 1 ml 1X PBS. Mounting medium was used with DAPI (4=,6-diamidino-2-phenylindole; Vector Laboratories Inc., catalog no. H-1200) to mount coverslips to slides. Slides were imaged by confocal microscopy on an LSM 700 laser scanning confocal microscope. The intensity levels of the images were adjusted such that no data were removed from images.

### Intracellular growth assays

HFFs grown on coverslips in 24 well plates were infected with freshly egressed tachyzoites pre-treated with either DMSO or 10μM hit compounds. and grown in confluent HFF coverslips for 24 h. Coverslips were fixed and stained with the anti-Toxo-FITC antibody as described above. Slides were imaged on an LSM 700 laser scanning confocal microscope. Cover slips were counted, and at least 150 vacuoles with 8 parasites/vacuole were counted.

### Extraction of Lipids from Parasites and TLC Analysis

For total lipids collection, parasites were grown in confluent HFF for 24 h. parasites were collected by centrifugation at 1000g and washed twice with 1X PBS. Equal number of parasites were collected, extracted with 1.5 ml of 2:1 chloroform/methanol by vortexing at room temperature for 1hr. 0.3 ml of ddH2O was added, the samples were vortexed on high for 1 minute, and phases were separated by centrifugation at 1000 rpm in a clinical centrifuge at room temperature. The upper phase was removed by aspiration and the organic phase washed with 0.25 ml 1:1 methanol/water. After phase separation, the lower organic phase was transferred to a new borosilicate tube and dried down under a stream of liquid nitrogen. Chloroform-resuspended samples were loaded onto silica gel TLC plates and resolved in 1D twice, first using chloroform/methanol/H2O (65:25:4) to mobilize the polar lipids and secondly using Hexane/acetone (100:1) to mobilize the polar lipids. For visualization of the separated lipids, the developed TLC plates were air dried and dipped uniformly in 8% (w/v) H3PO4 containing 10% (w/v) copper(II) sulfate pentahydrate, charred at 180 for 10min. The lipids were quantified using densitometry and imaged using scanner.

### Sample Preparation for Lipidomics

For lipid analysis, 5 T25 flask of confluent monolayer of HFF were infected with parasites and grown for 24 hrs. Parasites were subsequently collected and resuspended in 1.5 ml of 1xPBS buffer and transferred into glass vials pre-loaded with 2:1 chloroform/methanol/1XPBS to a final ratio of 2:1:1 chloroform/methanol/1xPBS.. Samples were vigorously shaken and lipid extraction was performed explained above. Five vials of lipids extracted from each strain (wild type and Δ*Tg*ASh4 parasites) were dried under nitrogen gas flow. Lipids were temporarily stored on dry ice before being transferred to a −80 freezer awaiting analysis on Q-TOF.

### Lipid analysis using high-performance liquid chromatography-mass spectrometry

Mass spectrometry analysis was performed with an electrospray ionization source on an Agilent 6545 Q-TOF LC/MS in positive and negative ionization modes. For Q-TOF acquisition parameters, the mass range was set from 100 to 1200 m/z with an acquisition rate of 10 spectra/second and time of 100 ms/spectrum. For Dual AJS ESI source parameters, the drying gas temperature was set to 250°C with a flow rate of 12 l/min, and the nebulizer pressure was 20 psi. The sheath gas temperature was set to 300°C with a flow rate of 12 l/min. The capillary voltage was set to 3500 V and the fragmentor voltage was set to 100 V. For separation of nonpolar metabolites, reversed-phase chromatography was performed with a Luna 5 mm C5 100 Å LC column (Phenomenex cat # 00B-4043-E0). Samples were injected at 20 ul each. Mobile phases for positive ionization mode acquisition were as follows: Buffer A, 95:5 water/methanol with 0.1% formic acid; Buffer B, 60:35:5 isopropanol/methanol/water with 0.1% formic acid. Mobile phases for negative ionization mode acquisition were as follows: Buffer A, 95:5 water/methanol with 0.1% ammonium hydroxide; Buffer B, 60:35:5 isopropanol/methanol/water with 0.1% ammonium hydroxide. All solvents were HPLC-grade. The flow rate for each run started with 0.5 minutes 95% A / 5% B at 0.6 ml/min, followed by a gradient starting at 95% A / 5% B changing linearly to 5% A / 95% B at 0.6 ml/min over the course of 19.5 minutes, followed by a hold at 5% A / 95% B at 0.6 ml/min for 8 minutes and a final 2 minute at 95% A / 5% B at 0.6 ml/min. Raw files were converted to mzXML format with MSConvert (ProteoWizard) using the Peak Picking Vendor algorithm. Files were analyzed using the web-based XCMS platform (Tautenhahn et al., 2012) with the following settings: signal to noise threshold, 6; maximum tolerated m/z deviation, 30 ppm; frame width for overlapping peaks, 0.01; and peak width, 10-60 s. Integrated peak intensities were normalized between conditions by median fold change. Ions were matched to the METLIN database with a 10 ppm tolerance for mass error. The tables containing potential metabolite annotations for each mass feature were downloaded from the XCMS platform. In order to analyze the metabolites according to their chemical classes, metabolite names were transformed into universally readable chemical identifiers: Using python 3 and the pubchempy wrapper for the PubChem PUG REST API (Fahy et al., 2007) metabolite names were searched against the PubChem database (Kim et al., 2018) and the PubChem CID and SMILES data were matched to the respective mass features. If a given mass feature could correspond to numerous metabolites, all potential metabolites were considered.

For annotation and classification of lipids, the PubChem CID’s were next searched against the LIPID MAPS^®^ Structure Database (LMSD) using their LIPID MAPS REST service using python 3 (Fahy et al., 2007). Finally, the resulting lipid annotations were matched to the original XCMS metabolomics data. Lipid annotations and their fold changes were plotted after grouping by “core” or “main class.” Plots were produced using python 3 with pandas, matplotlib, and seaborn packages. Boxes indicate the first, second (median), and third quartile ranges for fold change, while whiskers extend 1.5 times past the first or third quartile ranges. Values lying outside these ranges are indicated by circles.

### Library screening with purified proteins

For each covalent inhibitor, DMSO stocks were diluted in reaction buffer (20 mM HEPES, 150 mM NaCl, 10 mM CHAPS, pH 7.4) to a final concentration of 300 mM, followed by the addition of purified recombinant depalmitoylase at a final concentration of 50 nM. Reactions were incubated at 37°C and fluorescence was measured (Ex = 410 nm, Em = 450 nm) over 30 minutes. Data points collected from three replicates were normalized to the DMSO control and initial rates were calculated from a linear fit of the activity curves and expressed as relative fluorescence units per second (RFU/sec).

## Acknowledgments

We thank past and present Bogyo, Boothroyd, Egan, and Yeh laboratory members for their input and suggestions. This work was funded by NIH grant R01GM111703 (M.B.), B.M.B. was supported by the A. P. Giannini Foundation, M.L. was supported by Deutsche, Forschungsgemeinschaft (DFG) for funding under the Walter Benjamin Program

## Author contribution

O.O Designed and conducted the experiments, O.O. and M.B. conceptualized the work, analyzed data, and wrote the paper., O.O., B.M.B, and M.L analyzed Lipidomics data. M.N., K.L. P.C., and O.O. analyzed sequence, phylogenetic, and homology modeling. S.M.T and J.Z.L Lipidomics analysis by LC-MS. I.T.F. generated recombinant plasmid constructs. N.A synthesis QStE and QSE substrates.

