## Supplemental Figures for "*Toxoplasma gondii* serine hydrolases regulate parasite lipid mobilization during growth and replication within the host"

A

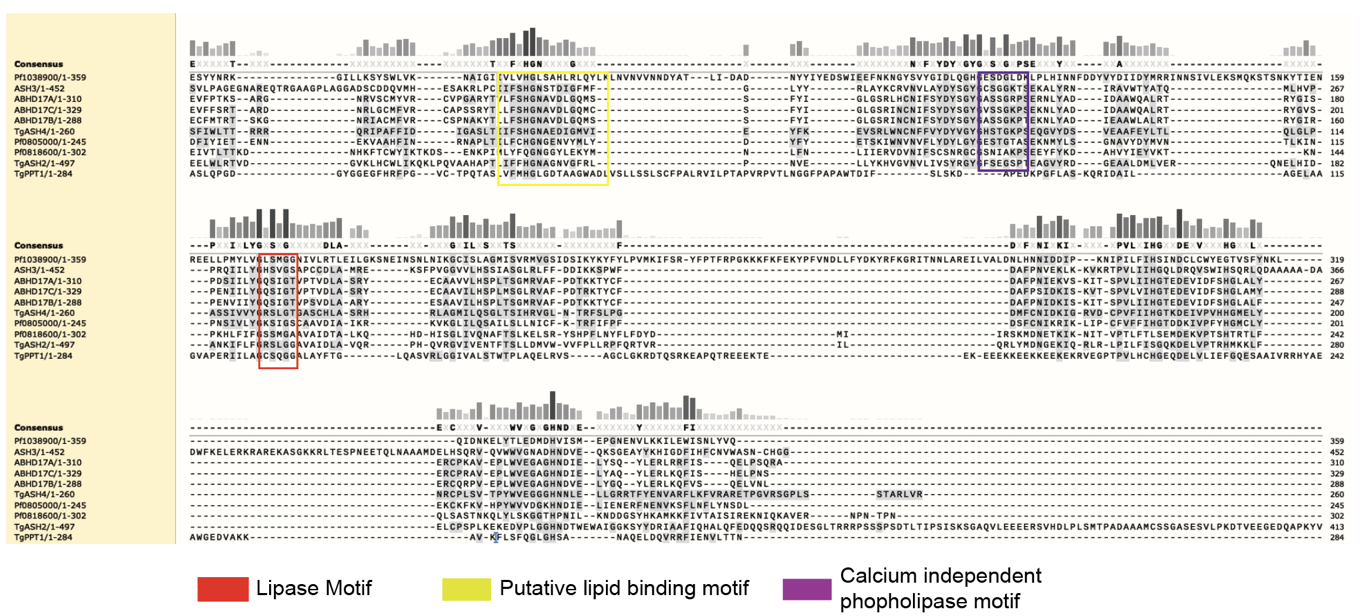

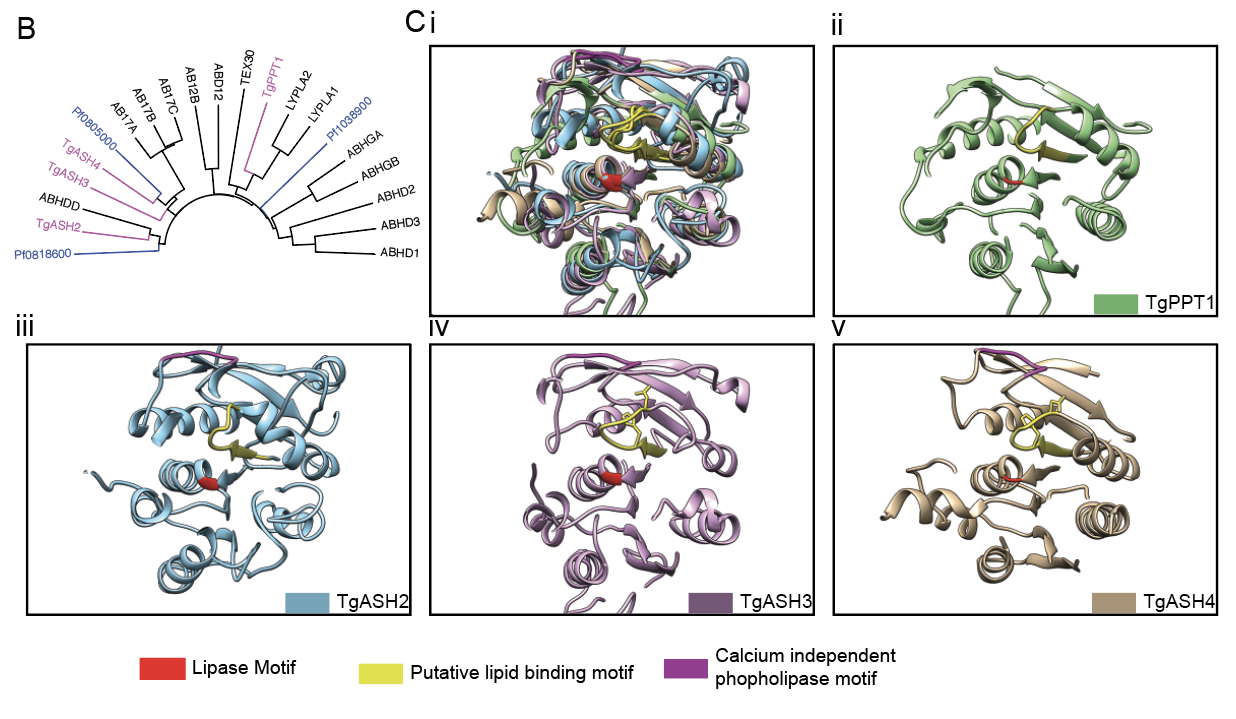

**Supplementary Figure 1. Sequence analysis of T. gondii and P. falciparum serine hydrolases (related to Figure 1)** (A) Sequence alignment of *Toxoplasma gondii* and *Plasmodium falciparum* serine hydrolases using Clustal Omega. Identical and highly similar residues are indicated within the conserved lipase motif (red), putative lipid binding domain (yellow) and calcium independent phospholipase motif (magenta). (B) Dendrogram of human, *T. gondii* and *P. falciparum* serine hydrolases showing the clustering in relation to human serine hydrolases (C). PyMOL structural modeling of *T. gondii* serine hydrolases using template structures (Table S1). The conserved lipase motif (red), putative lipid binding domain (yellow) and calcium independent phospholipase motif (purple) are highlighted. Images of (i) overlay of *Tg*PPT1 and *Tg*ASH2-4 (ii) *Tg*PPT1 (iii) *Tg*ASH2 (iv) *Tg*ASH3 and (v) *Tg*ASH4 homology models

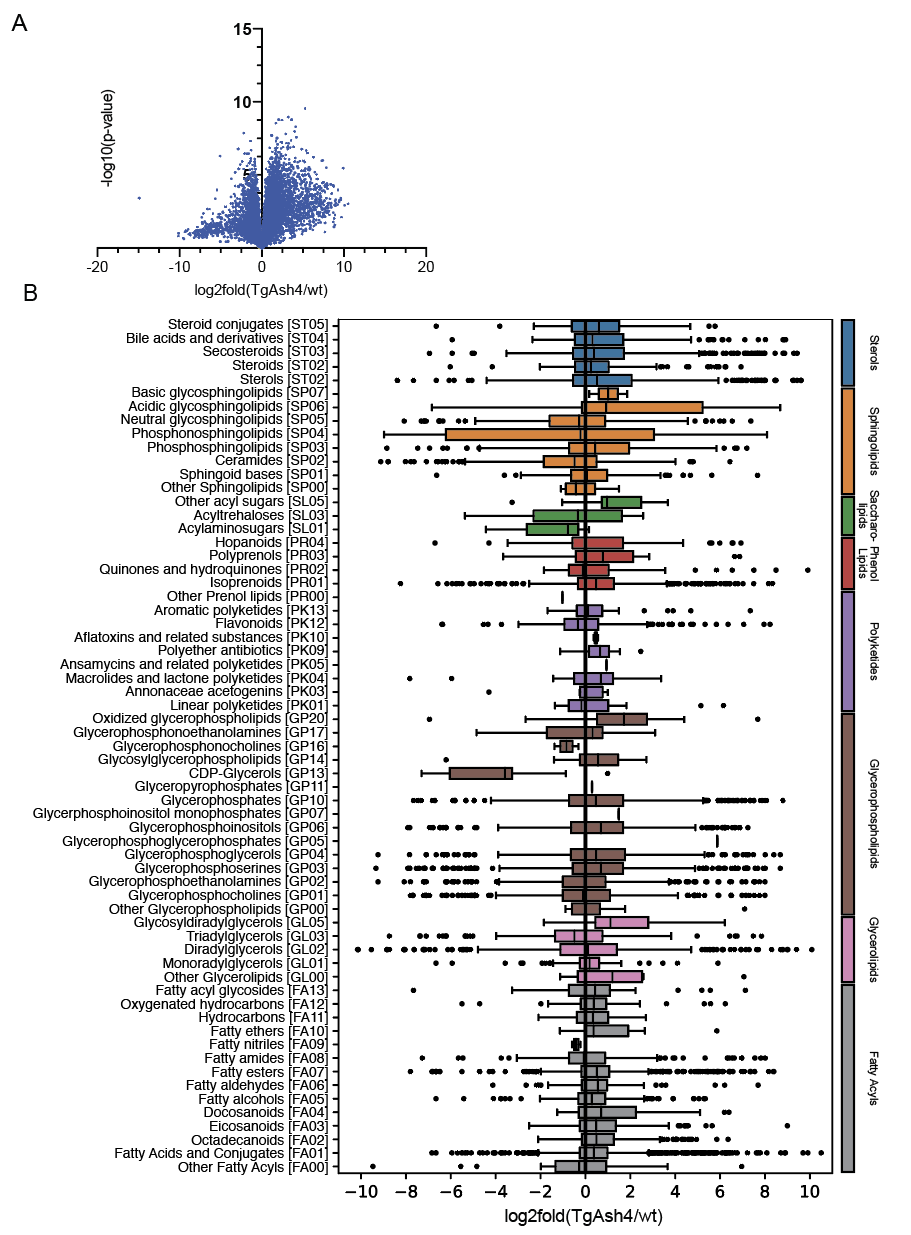

**Supplementary Figure 2. Lipidomic analysis of ΔTgAsh4 and wild type parasite (related to Figure 4)** Volcano plot showing log scaled mean fold change and adjusted p values for all the lipids resulting from the differential expression analysis between Δ*Tg*ASH4 and wild-type parasites (Figure 4C-E). (B) Box plot diagram highlighting the changes in the overall lipid species distribution in Δ*Tg*ASH4.

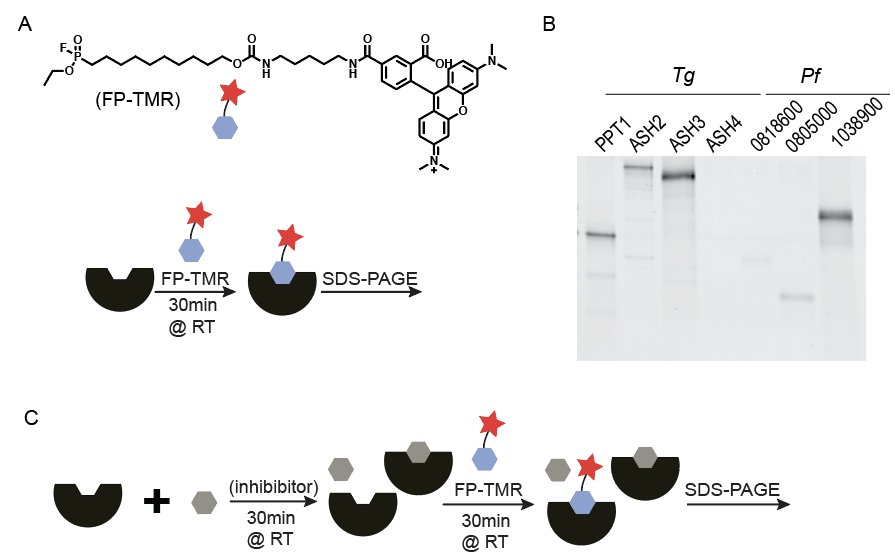

**Supplementary Figure 3. Active site labeling of recombinant serine hydrolases with Fp-TMR. Related to figure 5).** (A) Structure of FP-TAMRA (upper panel) and schematic showing labeling of the enzymes with FP-TAMRA before analysis of SDS_PAGE gel (lower panel) (B) SDS-PAGE gel showing the FP-TAMRA labeling of the recombinant *T. gondii* and *P. falciparum* serine hydrolases. (C) Schematic showing the inhibitor screen set using competition labeling with FP-TMR.

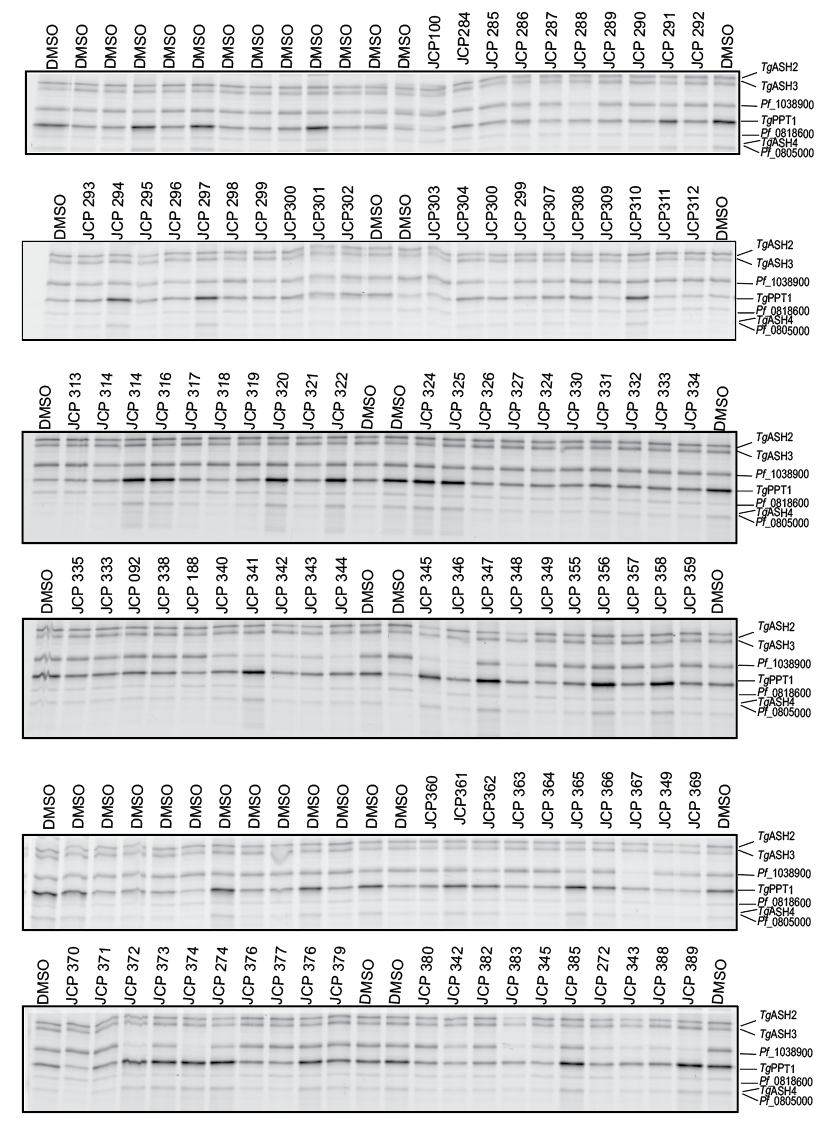

**Supplementary Figure 4.** SDS-PAGE gel based competitive activity-based protein profiling results showing inhibition of recombinant T. gondii and P. falciparum serine hydrolases. The enzymes were pooled and pre-incubated with 33µM of serine reactive compounds with and subsequent addition of FP-TAMRA. Competition was assessed using fluorescence intensity of SDS-PAGE gel bands.

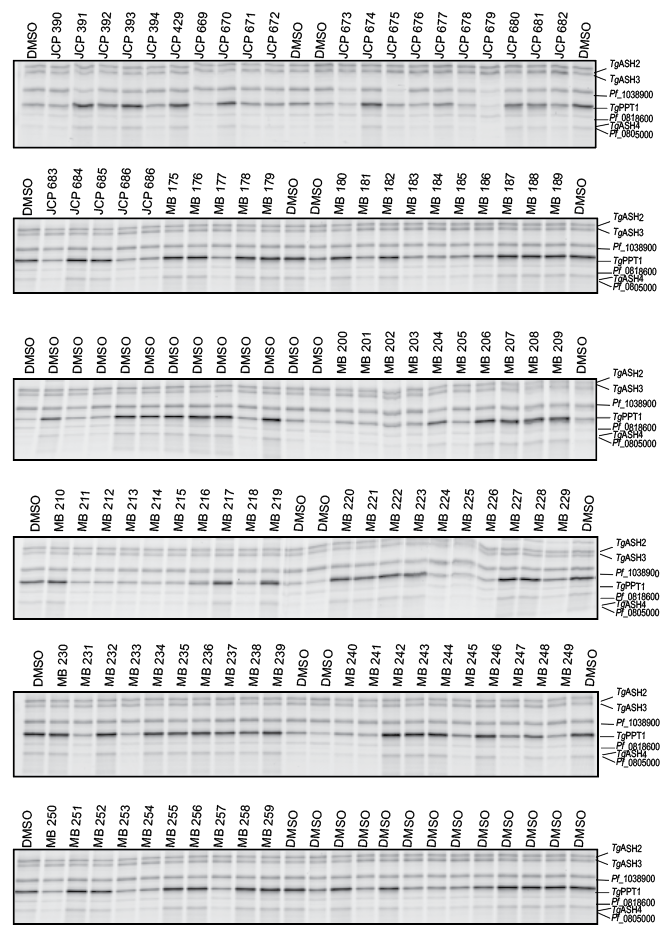

**Supplementary Figure 5.** SDS-PAGE gel based competitive activity based protein profiling results showing inhibition of recombinant *T.gondii* and *P. falciparum* serine hydrolases. The enzymes were pooled and pre-incubated with 33µM of serine reactive compounds and subsequent addition of FP-TAMRA. Competition was assessed using fluorescence intensity of SDS-PAGE gel bands.

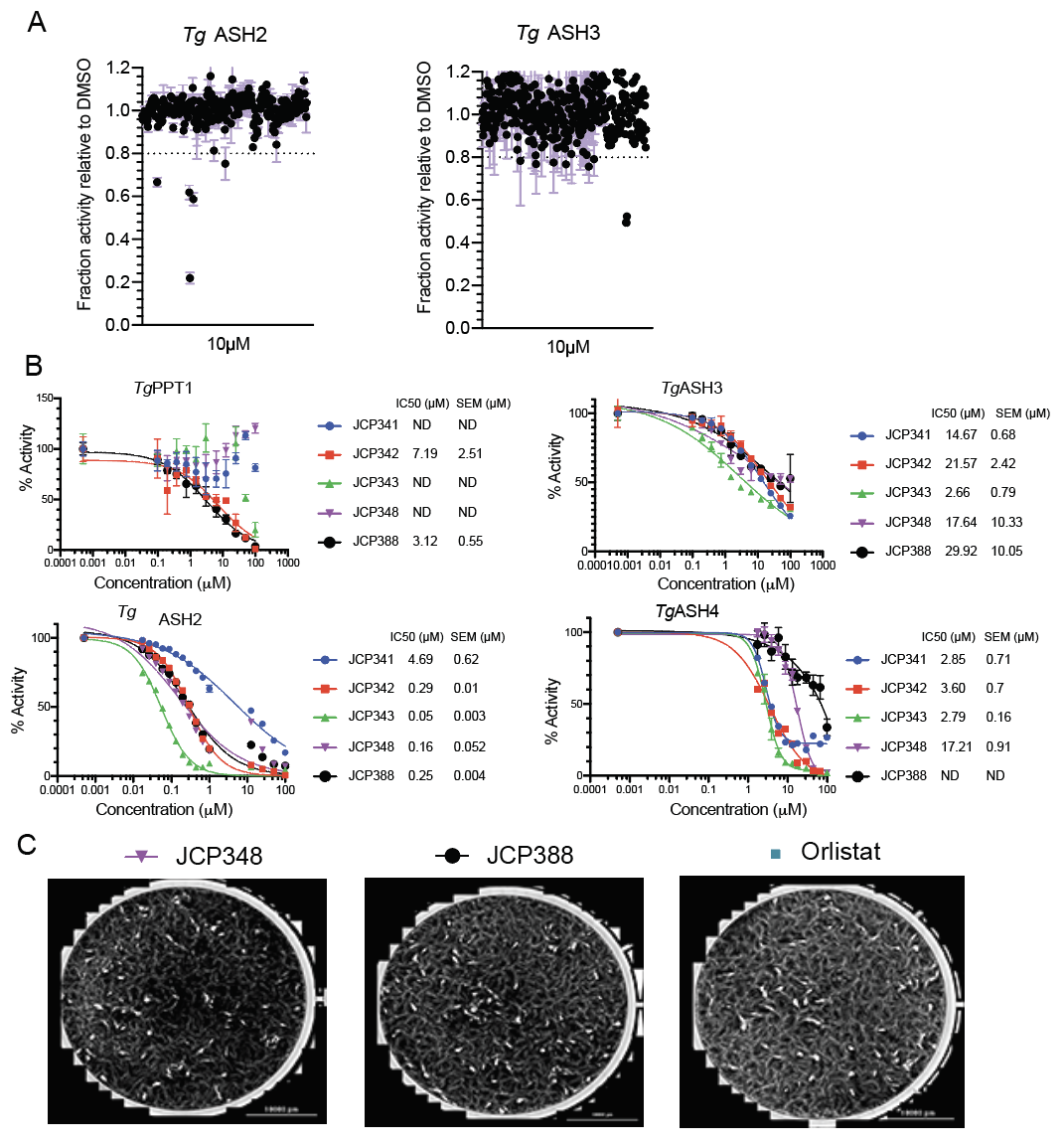

**Supplementary Figure 6. (**A) Dot plot showing our secondary screen inhibitors using a 4MU-octanote as the substrate (Figure 5B) and (B) Inhibitor validation in vitro and calculation of the IC50 curves and values of selected compounds (Figure 5B) (C) Inhibitor validation in cell culture infection model using a plaque assay that measures plaque formation by the wild type parasites in presence or absence of JCP341, JCP342, JCP343, JCP348, and Orlistat (Figure 6A, B).

**Table S1.** List of structural temples used to build homology models of serine hydroalses.

| **Enzyme** | **structural template** | **template type** | **sequence identity (%)** | **FFAS score** | **position of catalytic residues** | **remarks** |
| --- | --- | --- | --- | --- | --- | --- |
| *Tg*PPT1 |  |  |  |  |  |  |
|  | 5syn_A | thioesterase | 36 | -83.6 | S128, D221, H263 | analyzed model, position of docked substrate follow exper. position of inhibitor in the template structure** |
|  | 1fj2_A | thioesterase | 34 | -81.9 |  |  |
|  | 2wtm_A | EST1E | 15 | -26.8 |  | not well defined binding pocket |
| *Tg*ASH2 |  |  |  |  |  |  |
|  | 6imp_A | RTX toxin, serine hydrolase | 17 | -65 | S192, D268, H298 | analyzed model** |
|  | 5g59_A | esterase | 17 | -52.9 |  | second model** |
|  | 6ii2_A | putative RTX toxin | 15 | -31 |  | catalytic triad not formed |
| *Tg*ASH3 |  |  |  |  |  |  |
|  | 6imp_A | RTX toxin, serine hydrolase | 16 | -61.8 | S277, D346, H420 | H420 not modelled, template too short |
|  | 5g59_A | esterase | 15 | -47.2 |  | analyzed model** |
|  | 6ii2_A | putative RTX toxin | 15 | -29.9 |  | catalytic triad not formed |
| *Tg*ASH4 |  |  |  |  |  |  |
|  | 6imp_A | RTX toxin, serine hydrolase | 16 | -66.4 | S124, D188, H217 | position of D188 incorrect |
|  | 5g59_A | esterase | 20 | -58.8 |  | analyzed model** |
|  | 2wtm_A | EST1E | 18 | -37.6 |  | not well defined binding pocket |
